# Convergence of DNA methylation profiles in a novel environment in the reef coral *Porites astreoides*

**DOI:** 10.1101/747840

**Authors:** James L. Dimond, Steven B. Roberts

## Abstract

Phenotypic acclimatization is an organismal response to environmental change that may be rooted in epigenetic mechanisms. In reef building corals, organisms that are severely threatened by environmental change, some evidence suggests that DNA methylation is an environmentally responsive mediator of acclimatization. We investigated changes in DNA methylation of the reef coral *Porites astreoides* in response to simulated environmental change. Coral colonies were sampled from a variety of habitats on the Belize Barrier Reef and transplanted to a common garden for one year. We used restriction site associated DNA sequencing, including a methylation-sensitive variant, to subsample the genome and assess changes in DNA methylation levels after a year in the common garden. Methylation changes among the 629 CpG loci we recovered were subtle, yet coral methylomes were more similar to each other after a year in the common garden together, indicating convergence of methylation profiles in the common environment. Differentially methylated loci showed matches with both coding and non-coding RNA sequences with putative roles in intracellular signaling, apoptosis, gene regulation, and epigenetic crosstalk. There was a weak but positive and significant relationship between genetic and epigenetic variation, providing evidence of methylation heritability. Altogether, our results suggest that DNA methylation in *P. astreoides* is at least somewhat responsive to environmental change, reflective of the environment, and heritable, characteristics necessary for methylation to be implicated as part of potential transgenerational acclimatization responses.

## Introduction

Epigenetic processes, which contribute to gene regulation without affecting underlying DNA sequences, are increasingly recognized as molecular mechanisms that shape phenotypic responses (Duncan et al. 2014). Moreover, epigenetic signatures of organisms can change over their lifetimes, acting as potential records of, and responses to, environmental changes (Duncan et al. 2014; Hofmann 2017; Eirin-Lopez and Putnam 2019). The expanding evidence for both consequential functions of epigenetic processes and their plasticity is therefore driving interest among environmental and evolutionary biologists in search of the molecular basis of phenotypic plasticity, local adaptation, and acclimatization to climate change (Duncan et al. 2014; Hofmann 2017; Eirin-Lopez and Putnam 2019).

Although epigenetics encompasses a suite of molecular processes that appear to interact together, DNA methylation is the best understood and most widely studied of these processes (Eirin-Lopez and Putnam 2019). In the animal kingdom, 5-methylcytosine is the most common form of DNA methylation and is almost exclusively associated with CpG motifs. Invertebrate genomes are generally more sparsely methylated than vertebrate genomes, and methylation tends to be concentrated within gene bodies (i.e., introns and exons) of housekeeping genes (Sarda et al. 2012). While our understanding of the function of gene body methylation is only in its infancy, current evidence suggests that it helps ensure transcriptional fidelity, consistency, and efficiency, and may also be involved in alternative mRNA splicing (Neri et al. 2017; Flores et al. 2012).

Stable yet labile, epigenetic marks like DNA methylation can persist over generations, but they can also be primed and altered by environmental changes. In this way, epigenetic processes are thought to impart environmental “memories” in organisms (Iwasaki and Paszkowski 2014). Environmental memories may be particularly relevant in organisms such as plants and sessile invertebrates because they must weather any changes in their environment. Indeed, there are numerous examples of environmentally inducible epigenetic modifications in plants (Kinoshita and Seki 2014; Iwasaki and Paszkowski 2014). For sessile invertebrates, however, far less is known, and studies are just beginning to emerge.

Tropical reef corals are long-lived, sessile invertebrates that are thought to be particularly reliant on physiological acclimatization and phenotypic plasticity to cope with environmental variation (Gates and Edmunds 1999; Todd 2008). The underlying basis of this plasticity could lie, at least in part, in epigenetic mechanisms like DNA methylation (Roberts and Gavery 2012). However, in order to mediate phenotypic plasticity and acclimatization, DNA methylation itself must be plastic. To date, only a few studies have evaluated the response of DNA methylation to environmental change in corals. In a comparative study of two species of corals, Putnam et al. (2016) found that global methylation levels of *Montipora capitata* did not change in response to reduced pH conditions, while those of *Pocillopora damicornis* were responsive. In another study of simulated ocean acidification conditions with *Stylophora pistillata*, Liew et al. (2018b) observed modifications in methylation levels of genes involved in cell cycle and body size pathways that were reflected by phenotypic changes. Finally, in a reciprocal transplant study, Dixon et al. (2018) reported genome-wide changes in methylation in *Acropora millepora* that were correlated with physiological and transcriptional plasticity. In the latter two studies, there was evidence that changes in methylation were associated with acclimatization. However, we still lack sufficient information on the extent to which DNA methylation responds to environmental change.

In this study, we used a common garden transplantation approach to investigate the response of DNA methylation to environmental change in the Western Atlantic reef coral *Porites astreoides*. Corals were resampled after a year in a common garden and methylation levels were assessed using restriction site associated DNA sequencing (RADseq) techniques. We hypothesized that DNA methylation would be responsive to this manipulation and that methylation profiles among colonies would be more similar to each other after a year in a common environment.

## Methods

### Common garden experiment

In November 2015, 19 colonies of *P. astreoides* (approximately 20 cm diameter) were transplanted from their home site to a common garden in the shallow (∼1 m depth) backreef in front of Carrie Bow Cay (CBC; 16° 48’ 9′′N, 88° 4’ 55′′W), Belize. Coral colonies were haphazardly selected from shallow habitats (1-3 m depth) within a 20 km radius of CBC. Some colonies were collected from windward backreef habitats similar to those of CBC, while others were collected from inshore habitats. Upon collection, colonies were first sampled for DNA by chipping off a small piece of the colony with hammer and chisel, then preserving the fragment in salt-saturated DMSO solution at room temperature until extraction. Colonies were then halved, and one half was brought to the common garden, an approximately 9 m^2^ area, where they were reattached to the substratum using A-788 splash zone compound. Fourteen of the original 19 colonies also had their remaining half reattached to the substratum from which they were collected, serving as controls. Numbered aluminum tags were attached adjacent to each coral to permit later identification, and subsurface floats were moored near all control colonies left at their site of origin. Colonies were resampled one year later in November 2016, again removing and preserving a small fragment for DNA extraction.

### Environmental analysis

To characterize the surrounding physical environment, remotely sensed data from the AQUA MODIS satellite sensor were used (NASA 2016). Previous studies have reported strong correlations between remotely sensed sea surface temperatures (SST) and those recorded from data loggers moored in shallow benthic habitats, suggesting that remotely sensed data is appropriate for studies of shallow benthic environments (Smale and Wernberg 2009; Pearce et al. 2006). Monthly AQUA MODIS climatology datasets for the period November 2015 to October 2016 were used. For SST, we used the 11μ band nighttime dataset (NASA 2016). For chlorophyll concentration, the OCx algorithm was used (NASA 2016). Datasets were imported into R as raster images for analysis. Additionally, Belize basemap (Meerman and Clabaugh 2017) and coral reef basemap (UNEP-WCMC 2018) shapefiles were used to provide spatial context.

### DNA extraction and sequencing

DNA extraction and library preparation and sequencing followed methods described in detail by Dimond et al. (2017). Briefly, libraries were prepared according to the double-digest RADseq (ddRADseq) and EpiRADseq methods of Peterson et al. (2012) and Schield et al. (2016), respectively. This created tandem libraries for each sample; the ddRADseq library used a methylation insensitive common cutter (MspI) targeting 5’-CCGG-3’ motifs, while the EpiRADseq library used a methylation sensitive common cutter (HpaII, also targeting 5’-CCGG-3’). ddRADseq and EpiRADseq rely on a size-selection step to ensure targeted sequencing of a small subset of fragments within a narrow size range, with the ultimate goal of sequencing genomic intervals that will be present across many samples (Peterson et al. 2012). Paired-end, 100 bp libraries were sequenced in equimolar ratios on the Illumina HiSeq 4000. A total of 96 samples, half ddRADseq and half EpiRADseq, were sequenced on a single lane; some of these samples were used in a separate study.

### Symbiont genotyping

Most reef corals engage in obligate symbiotic associations with dinoflagellates of the family Symbiodiniaceae (LaJeunesse et al. 2018). To inform the sequence assembly workflow described below regarding the choice of symbiont genome used to subtract symbiont reads, symbiont genotypes for each sample were determined at the genus (formerly clade) level via NCBI BLAST (v. 2.6.0) queries of ddRADseq libraries. A custom BLAST database was generated by searching for all *Symbiodinium* (formerly *Symbiodinium* clade A), *Breviolum* (*Symbiodinium* clade B), *Cladocopium* (*Symbiodinium* clade C), and *Durusdinium* (*Symbiodinium* clade D) records in the NCBI nucleotide database. Search terms were as follows: (((“*Symbiodinium*”[Organism] AND “*Symbiodinium* sp. clade A”[Organism]) OR “*Symbiodinium* sp. clade B”[Organism]) OR “*Symbiodinium* sp. clade C”[Organism]) OR “*Symbiodinium* sp. clade D”[Organism]. Due to wide variation in the number of records for each taxon, the full dataset was standardized by random sampling to retain 20,000 records per taxon (80,000 total records retained out of the original 447,000). The resulting sequences were used as a BLAST database against which each ddRADseq library was queried with BLASTN using max_target_seqs = 1 and an e-value of 1e-20. Reads that aligned to more than one taxon were removed, and any reads mapping to 18S rDNA were also removed since this gave many ambiguous matches.

The efficacy of this approach was tested using a prior dataset for which symbiont genotyping had also been performed using PCR amplification and Sanger sequencing of cp23S rDNA amplicons (methods described and data reported in Dimond et al. 2017). Ten samples of branching *Porites* spp. were tested for correspondence between symbiont identities determined via BLAST searches of ddRADseq reads to those determined via cp23S amplicon sequencing.

### Coral sequence assembly

Coral ddRADseq and EpiRADseq sequences were assembled with *ipyrad* v.0.5.15 (Eaton 2014), using the “denovo – reference” assembly method to exclude symbiont reads from the assembly. Given the results of the symbiont genotyping analysis identifying the dominant symbiont taxon as *Symbiodinium* (formerly *Symbiodinium* clade A; see Results) in all corals, the *Symbiodinium microadriaticum* (GenBank Accession no. GCA_001939145.1; Aranda et al. 2016) genome was used as a reference to exclude symbiont reads. Step one of *ipyrad* demultiplexed the reads by identifying restriction overhangs and barcode sequences associated with each sample; zero barcode mismatches were tolerated. Demultiplexed samples were then combined in a single directory for further steps. In step two, reads were trimmed of barcodes and adapters and quality filtered using a q-score threshold of 20, with bases below this score converted to Ns and any reads with more than 5 Ns excluded. Next, reads were mapped to the symbiont reference genome with bwa mem using default settings and any mapped reads were excluded from further analysis. Similar clusters of the remaining reads were then aligned using a threshold of 90% similarity. Step four performed joint estimation of heterozygosity and error rate (Lynch 2008) based on a diploid model assuming a maximum of 2 consensus alleles per individual. Step five used the parameters from step four to determine consensus bases calls for each allele and removed consensus sequences with > 5 Ns per end of paired-end reads. With consensus sequences identified, step six clustered and aligned reads for each sample to consensus sequences. Lastly, the data were filtered according to maximum number of indels allowed per read end (8 indels), maximum number of SNPs per locus (20), maximum proportion of shared heterozygous sites per locus (0.3) and minimum number of samples for a locus to be reported (20). From this final dataset, which included monomorphic loci, a subset of the data was chosen that maximized the number of resampled individuals (sampled in both 2015 and 2016) with a robust set of shared loci (i.e., loci with missing data were excluded). Several samples were excluded from analysis if either one or both of the years in which they were sampled had low data yield (e.g., due to low starting DNA quality). As a definition of terms used here, the term locus to refers to a consensus paired-end read. The term SNP refers to a single nucleotide polymorphism on a locus, while the term CpG refers to a cytosine-guanine dinucleotide pair that can be either methylated or non-methylated at the 5’-CCGG-3’ restriction site of each locus.

### Genetic analysis

Unlinked SNPs were scored by *ipyrad* at 1 SNP per locus with the least amount of missing data; SNPs were sampled randomly if they had equal amounts of missing data. The SNP error rate, defined as the proportion of SNP mismatches between pairs of datasets from identical individuals (Mastretta-Yanes et al. 2015), was estimated by treating ddRADseq and EpiRADseq libraries as technical replicates and calculating pairwise differences between individuals using the *dist.gene* function in the R package *ape* (Paradis et al. 2004).

### Epigenetic analysis

Analysis of DNA methylation using EpiRADseq data relies on read count information, as read counts of loci using this technique are inversely related to their methylation frequency (Schield et al. 2016). The analysis followed methods described in detail by Dimond et al. (2017) using a tandem ddRAD/EpiRAD approach in which reads that were present in the ddRAD library but absent in the EpiRAD library were considered methylated. Briefly, read counts for each locus obtained from *ipyrad* output were standardized according to library size for each sample using the R package *edgeR* (Robinson et al. 2010). As in Dimond et al. (2017), the residuals from linear regression of ddRAD vs. EpiRAD read counts were then used to ascertain methylation status, however, instead of manually setting the methylated/unmethylated threshold for a locus, k-means clustering was used to differentiate methylated from unmethylated loci using k=2. The *superheat* R package was used for heatmap visualization of methylation patterns (Barter and Yu 2018). The potential functions of differentially methylated loci were determined via a web-based BLASTN search using the default ‘nr’ nucleotide database and an e-value threshold of e^-5^.

### Relationships between genetic and epigenetic variation

Pairwise genetic and epigenetic distance were computed using the *dist.gene* function in the R package *ape* (Paradis et al. 2004). Comparisons between different fragments of the same individual (experimentally generated clones) were excluded. Linear regression analysis was used to test the hypothesis that pairwise genetic and epigenetic variation were positively related.

### Data accessibility

Demultiplexed sequence reads can be accessed in the NCBI Sequence Read Archive under accession number SRP132538 (https://trace.ncbi.nlm.nih.gov/Traces/sra/?study=SRP132538), or under BioProject PRJNA433592 (https://www.ncbi.nlm.nih.gov//bioproject/PRJNA433592). Analysis workflows can be accessed at https://github.com/jldimond/P.ast-transplant.

## Results

### Sample recovery and data yield

Of the 19 colony halves transplanted to the common garden, only one was not recovered a year later. Of the 14 control halves left at their site of origin, three were not relocated, while a fourth had been overtaken and killed by a damselfish garden. Although many samples were submitted for sequencing, low sequencing yield from several samples effectively excluded them from the final analysis due to insufficient data. Sequencing yield was highly variable, producing an average of 3.4 (s.d. = 3.0) million reads per sample. An average of 7539 (s.d. = 4799) consensus loci per sample were included in the final assembly. If data were insufficient (based on summary statistics given by *ipyrad*) for one or both sampling years for a given coral colony, the colony was excluded from analysis. The optimal balance between colony sample size and number of shared loci was achieved with N = 8 colonies sampled in both 2015 and 2016, including two controls, sharing 629 loci. Thus, 629 SNPs and CpGs shared across all samples were analyzed.

### Environmental analysis

Of the eight colonies included in the final analysis, four (colonies 5, 6, 10, and 11) originated from inshore environments approximately 6-8 km west of the barrier reef, while the other four (colonies 2, 3, 8, and 9) were collected from offshore environments just shoreward (within 500 m) of the barrier reef (Fig. 1). Inshore reefs were characterized by slightly cooler annual temperatures, a larger seasonal temperature range, and higher chlorophyll a concentration (Fig. 1A-C). These effects likely reflect the relatively shallow depth and longer residence time of water in the lagoon and its greater susceptibility to seasonal variation in solar heating and wind patterns, in addition to freshwater runoff and terrestrial nutrient sources. Most corals brought to the common garden originated in habitats with a greater seasonal temperature range and high chlorophyll concentration; the slightly lower mean temperature of these habitats was probably not biologically meaningful (Fig. 1D).

**Figure 1.**
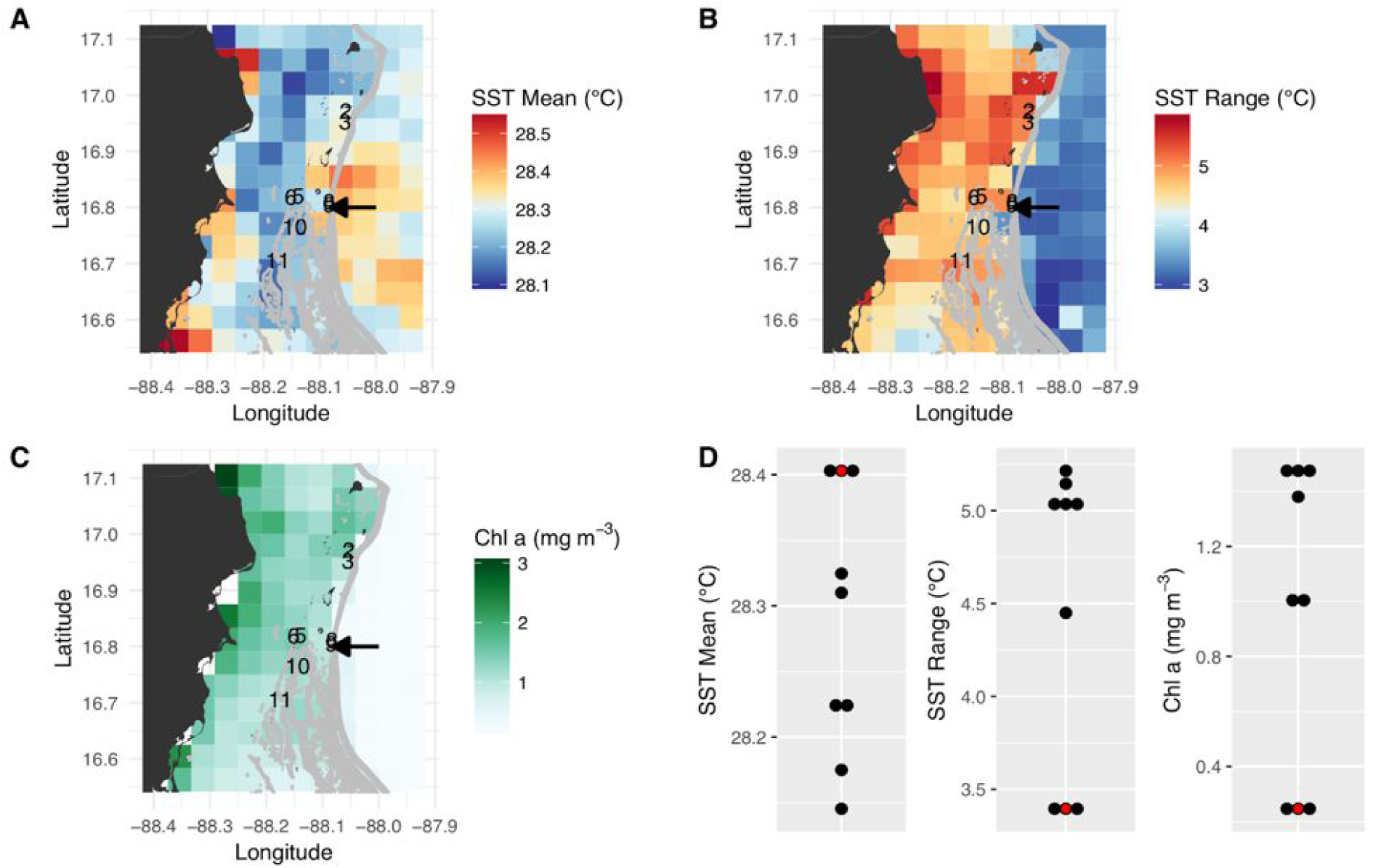
Satellite-derived environmental conditions associated with coral collection sites and the common garden along the Belize Barrier Reef. (A-C) Mean annual sea surface temperature (SST), annual SST range, and annual mean chlorophyll a concentration. The collection sites of the eight analyzed coral colonies are depicted by colony numbers, and the arrows show the location of the common garden (colonies 8 and 9 originated from just north of the common garden site). (D) Summary of native habitat and common garden environmental conditions. The common garden is depicted in red.

### Symbiont genotyping

For the branching *Porites* spp. test dataset evaluating the efficacy of symbiont genotyping via BLAST searches using ddRADseq reads, the dominant symbiont taxon detected via BLAST search was identical to the dominant symbiont taxon detected via cp23S Sanger sequencing in all cases (Fig. 2). Given these results, we proceeded with the BLAST approach with ddRADseq reads from *P. astreoides.* Among the *P. astreoides* samples, all individuals hosted ≥ 98% *Symbiodinium* (formerly *Symbiodinium* clade A) across both years. This justified the use of the *S. microadriaticum* genome to subtract symbiont sequences during de novo assembly of the RADseq data.

**Figure 2.**
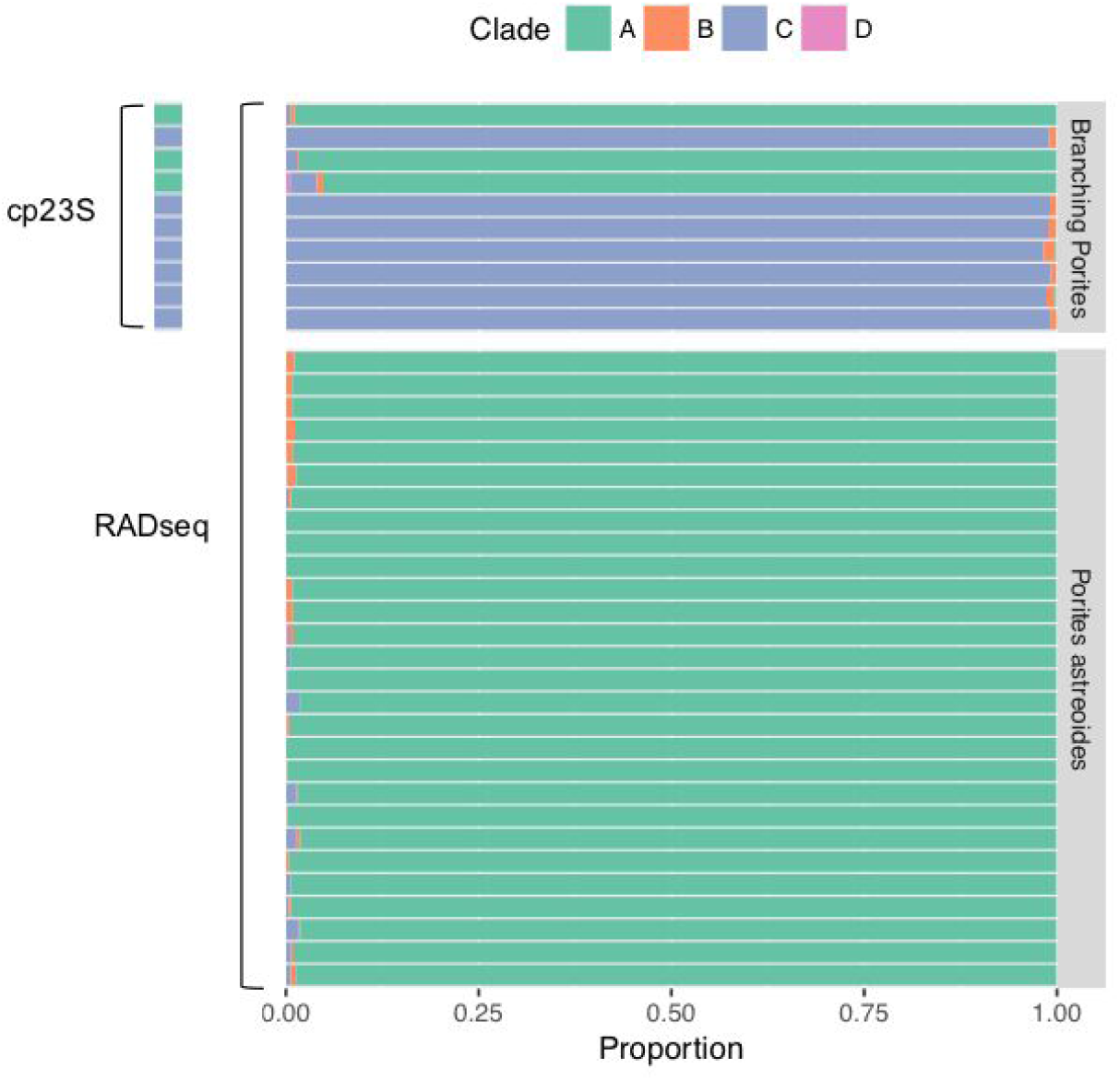
Symbiont identification using BLAST searches of ddRADseq data against custom databases of *Symbiodinium* (formerly *Symbiodinium* clade A), *Breviolum* (*Symbiodinium* clade B), *Cladocopium* (*Symbiodinium* clade C), and *Durusdinium* (*Symbiodinium* clade D). Top panel shows symbionts of branching *Porites* spp. collected for a prior study (Dimond et al. 2017); the dominant symbiont found in BLAST searches was identical to the dominant symbiont identified via cp23S Sanger sequencing. Bottom panel shows results from analysis of the *P. astreoides* colonies that are the primary subject of this study.

### Genetic analysis

Based on genotyping and analysis of genetic distance among technical replicates (ddRAD and EpiRAD libraries combined), the SNP error rate was estimated to be 1.2% (s.d. = 0.7%), well below previously published estimates of reduced representation sequencing data (Fig. 3; Mastretta-Yanes et al. 2015; Recknagel et al. 2015; Dimond et al. 2017). In other words, genotyping was 98.8% accurate. Genetic distance analysis also indicated that resampling of corals from 2015 to 2016 was accurate and no errors (e.g., sampling, misidentification) were made, showing SNP calling error comparable to the SNP error rate reported above for technical replicates (1.7%, s.d. = 0.9%). Chimerism (within-colony genetic variation resulting from fusion of juvenile colonies) and mosaicism (within-colony genetic variation arising from somatic mutations) is also not uncommon among scleractinian corals (Schweinsberg et al. 2015), and this analysis shows no evidence for these phenomena among the colonies sampled here. By contrast, genetic distance among all non-replicate and non-resampling pairwise comparisons between individuals was much greater, averaging 16.3% (s.d. = 1.4%). No clones were identified.

**Figure 3.**
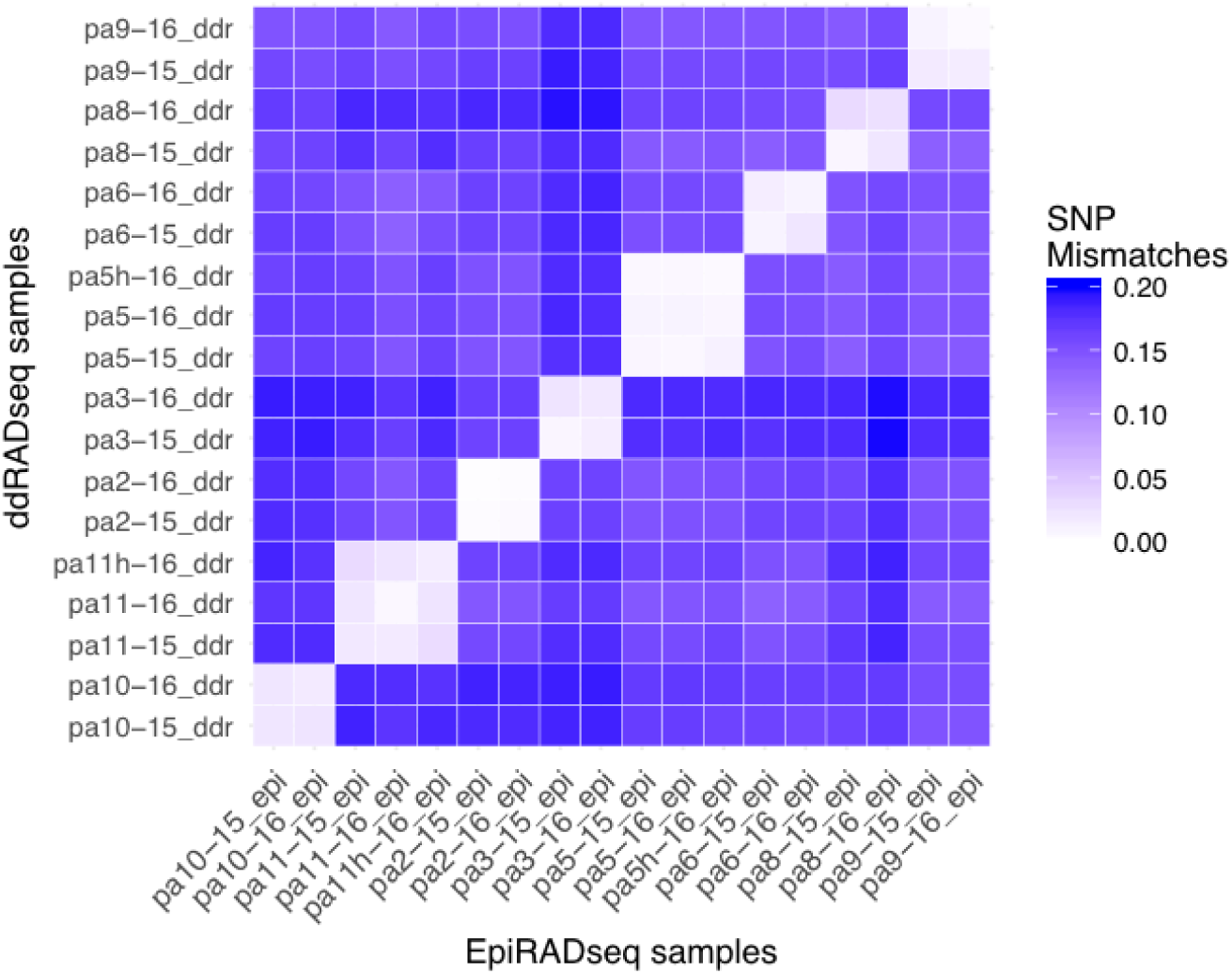
Pairwise SNP mismatches between the *P. astreoides* samples. EpiRADseq samples are shown along the x-axis, while ddRADseq samples are shown along the y-axis. Values along the diagonal illustrate the SNP error rate using the two libraries as technical replicates, while off-diagonal values show the error associated with resampling corals from 2015 to 2016. Two controls that were left at their site of origin (pa11h and pa5h) are included.

### Epigenetic analysis

Across all samples and years, an average of 18.6% (s.d. = 0.9%) of CpGs were methylated (Fig. 4). As indicated by the low variance, most loci were either methylated or unmethylated across all samples and years; 73% of loci were constitutively unmethylated across samples and years, 12% were constitutively methylated, and 15% were differentially methylated (Fig. 4). The low SNP error rate reported above also adds confidence to the epigenetic analysis as it indicates that reads were correctly assigned to consensus loci at the level of each sample.

**Figure 4.**
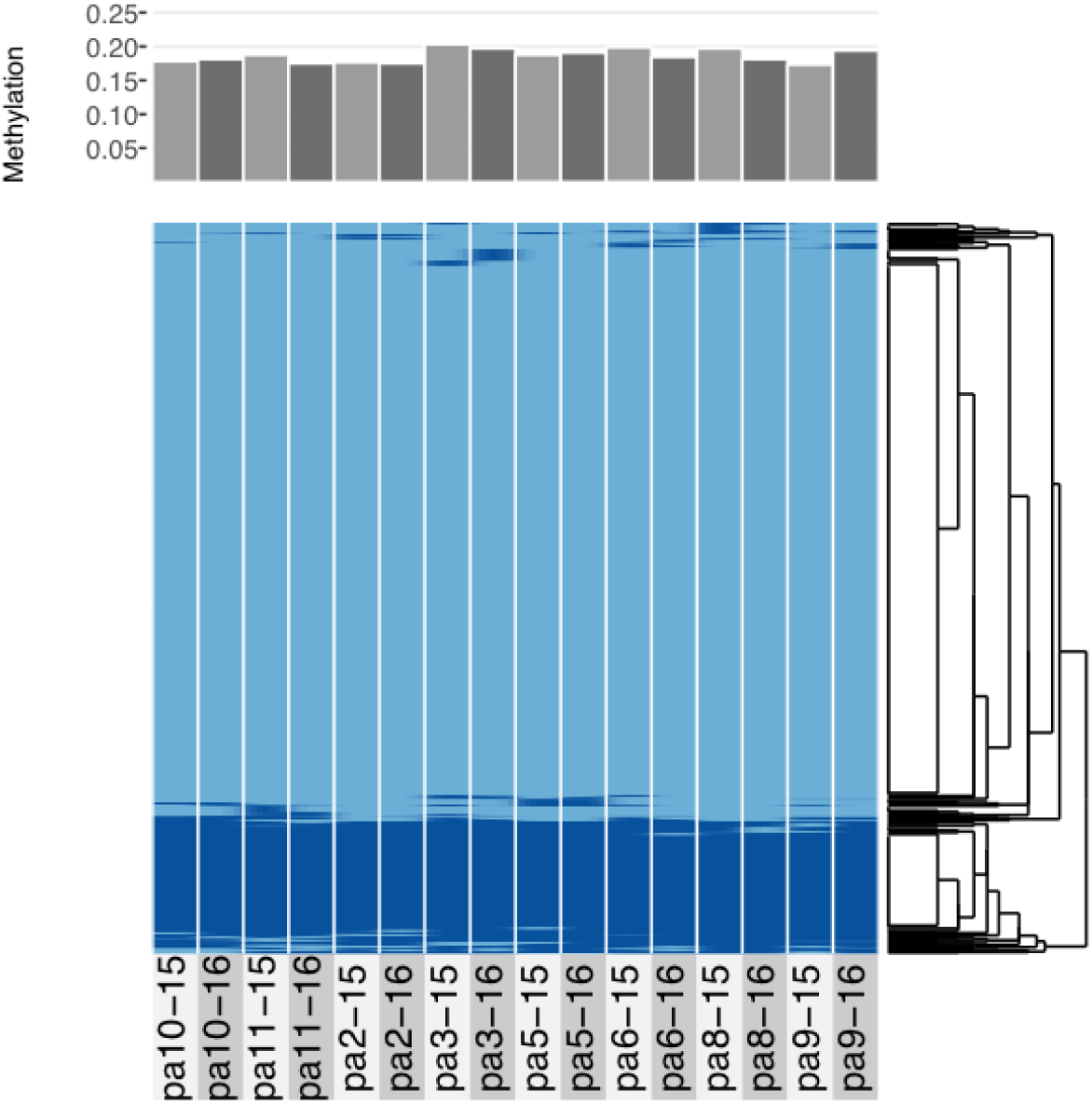
Comparative methylation profiles of the eight *P. astreoides* samples from 2015 to 2016. Every two vertical bars is a single individual, with the first bar showing 2015 methylation and the second showing 2016 methylation. Top panel shows proportion of methylated loci, while bottom panel shows a heatmap with methylated loci in dark blue and unmethylated loci in light blue, ordered via hierarchical clustering.

All corals underwent some degree of change in their methylation status, including the two controls (Fig. 5A). The mean percentage of loci changing methylation state per colony was 2.0% (s.d. = 0.9%). Among the two colonies with controls, the percentage of loci that changed methylation state was equivalent for colony pa11 and its control (2.9%), while for colony pa5, percent change was 1.0% and 1.7% for the common garden and control, respectively. Although there was no significant overall change in percent CpG methylation from 2015 to 2016 (paired t-test, df = 7, p = 0.511; Fig. 5), there was evidence for convergence of the methylation status among colonies toward a more similar methylome after a year in the common garden together, as pairwise differences between colonies were significantly lower in 2016 (paired t-test, df = 27, p < 0.001; Fig. 5).

**Figure 5.**
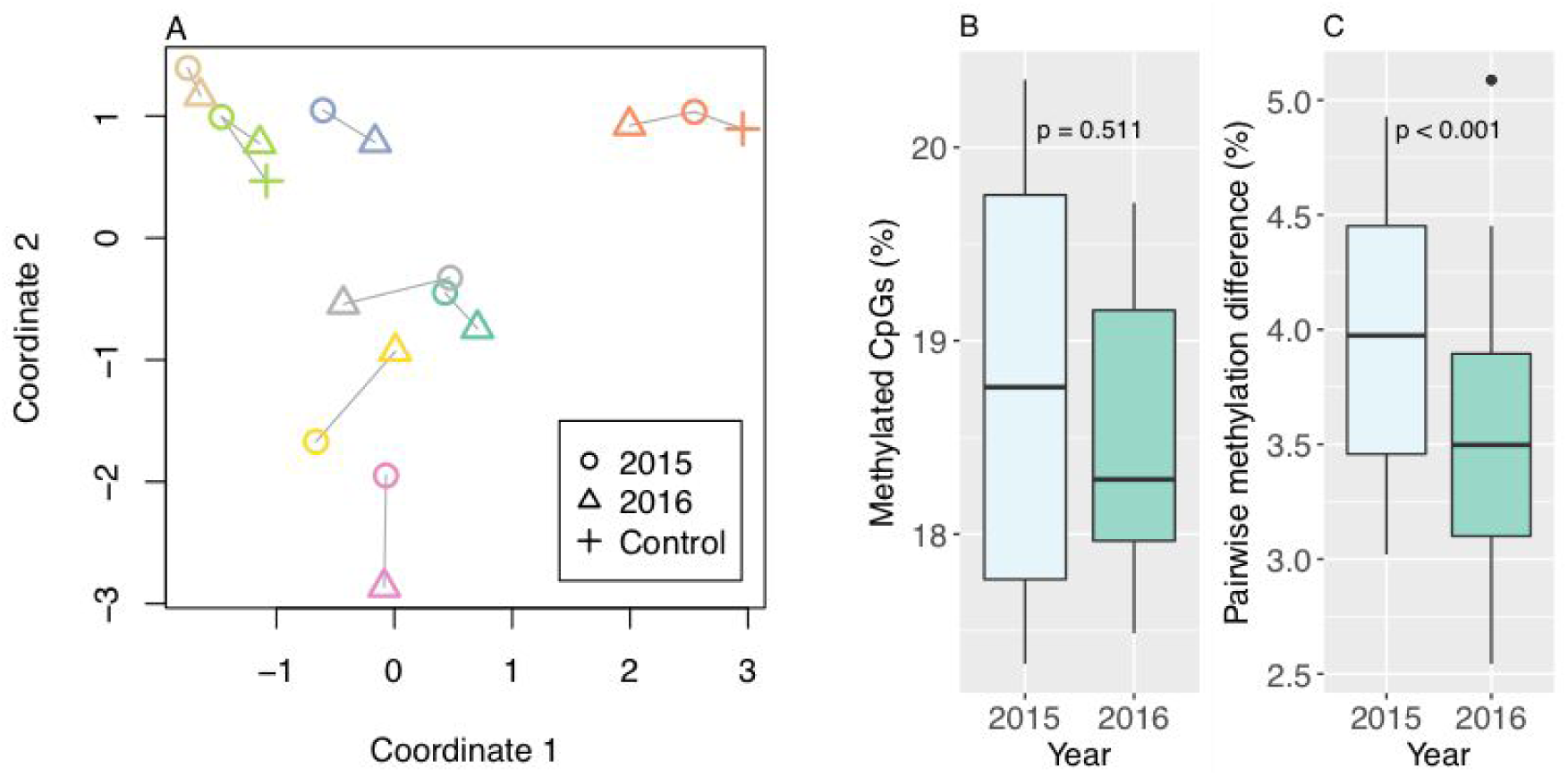
Methylation changes from 2015 to 2016. (A) Multidimensional scaling plot of methylation. Individual colonies are shown with different colors, and vectors show relative direction and magnitude of change from 2015 to 2016, including two controls that were left at their site of origin. (B) Percent methylated CpGs by year. (A) Pairwise methylation difference by year, including all pairwise comparisons between colonies.

Of the differentially methylated loci, most (76 of 95 = 80%) were differentially methylated only in a single individual colony. To further evaluate differentially methylated loci with some degree of confidence, only loci differentially methylated in more than one individual were considered further. Nineteen loci (19 of 95 = 20%) met this criterion and eight returned BLASTN hits. All matches were coral RNA sequences; six were mRNA sequences, while two were non-coding (nc) RNA sequences (Table 1). Four of the sequences had predicted products, including a protein kinase, a helicase-like ribonucleoprotein, a FAM98A-like protein, and a tax-1 binding protein homolog.

**Table 1.**
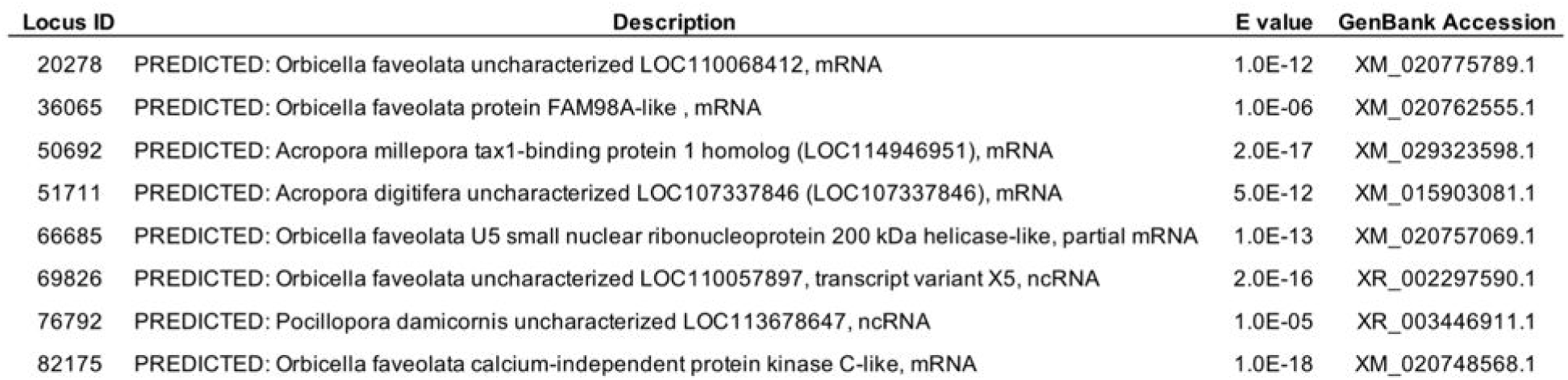
BLASTN hits for loci that were differentially methylated in at least 2 samples.

### Relationships between genetic and epigenetic variation

There was a weak yet significant and positive relationship between pairwise genetic and epigenetic distance (R^2^ = 0.131, p <0.001), indicating that corals that are more similar genetically also tend to be more similar epigenetically (Fig. 6). Genetic distance among colonies was also considerably greater than epigenetic distance (Fig. 6).

**Figure 6.**
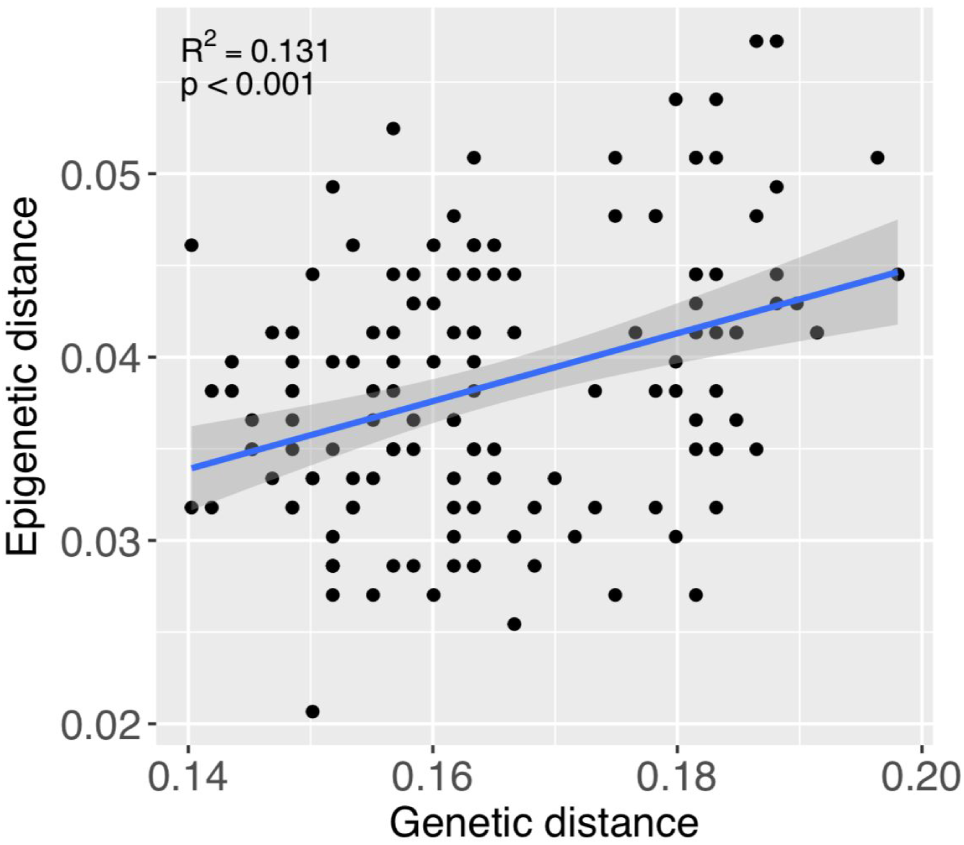
Relationship between pairwise genetic and epigenetic distance. Each point represents a pairwise comparison between each combination of samples, excluding comparisons between fragments of the same individuals. Coefficients in upper left region of figure were obtained from linear regression analysis. The trendline from this analysis is shown, with the shaded region around the line representing 95% confidence limits.

## Discussion

Our study shows evidence that DNA methylation in *P. astreoides* is at least somewhat responsive to environmental change, reflective of the environment, and heritable. These characteristics of methylation are necessary for it to be implicated as part of potential transgenerational acclimatization responses (Eirin-Lopez and Putnam 2019). Relatively fast acclimatization responses, along with slower natural selection, are essential, complementary processes that enable coral persistence in the face of climate change (Palumbi et al. 2014).

Transplantation to a new environment for a one-year period elicited subtle changes in the methylome of *P. astreoides*. Yet, there was also evidence that corals converged toward similar methylomes in the common garden, suggesting that the subtle changes in methylation were reflective of the new environment. Similarly, in a 3-month reciprocal transplant study of *Acropora millepora* on the Great Barrier Reef, Dixon et al. (2018) found that while DNA methylation was considerably less responsive than gene expression to transplantation, the methylomes of transplants became more similar to native corals. Moreover, methylation changes were correlated with measures of physiological competency in novel environments, with transplanted individuals whose methylomes became more similar to those of native individuals showing more robust physiological profiles (Dixon et al. 2018). The small yet significant convergence of methylation patterns observed after one year in the common garden in our study is in agreement with the results of Dixon et al. (2018).

Responsiveness of DNA methylation to changing environmental conditions is not a foregone conclusion, as shown by Putnam et al. (2016) in their study of global methylation responses of two coral species to low pH. The relatively environmentally sensitive coral *Pocillipora damicornis* exhibited significant changes in methylation, while the more robust coral *Montipora capitata* did not. Interestingly, whereas *P. damicornis* performed poorly and showed limited evidence of acclimatization to the 6-week exposure to experimental conditions, *M. capitata* performed well. This suggests that changes in methylation are not necessarily associated with acclimatization. However, bulk changes in methylated DNA were quantified by Putnam et al. (2016), so it is possible that individual genes in *M. capitata* underwent both increases and decreases in methylation to contribute to homeostasis. Indeed, this is the scenario reported by Dixon et al. (2018) and in our study. Here, approximately 18.5% CpG methylation was maintained before and after transplantation despite an average of 2% of loci changing their methylation state, reflecting a combination of increases and decreases in methylation among individual colonies and loci. Likewise, Dixon et al. (2018) reported no genome-wide increases or decreases in methylation in response to transplantation, but instead that methylation changed in a so-called “seesaw” pattern whereby changes in methylation among hypomethylated genes were mirrored by changes in the opposite direction among hypermethylated genes. Their study, along with a study by Liew et al. (2018b) on pCO_2_-mediated changes in methylation and phenotype in *Stylophora pistillata*, have concluded that environmentally induced changes in DNA methylation are associated with homeostatic regulation. The precise mechanisms of this regulation, however, will require further study to fully understand.

The relatively small changes in methylation we detected here could be at least partially attributed to the limited difference in native habitat environmental conditions relative to the common garden. Mean temperature among collections sites and the common garden varied only slightly, while temperature range and chlorophyll concentration were somewhat more variable. It is noteworthy, however, that Dixon et al. (2018) measured relatively small changes in methylation despite much greater environmental differences between transplantation habitats, albeit with only a three-month experimental duration.

Were methylation changes truly reflective of common garden conditions? This is worth considering since changes in methylation among the two control colonies that remained at their site of origin were similar in magnitude to the experimental colonies. This could suggest that methylation simply changes temporally as corals age, or that the experimental act of transplantation (halving colonies and reattaching them) caused methylation changes. Age-related changes in methylation, for example, are well-documented among humans and other vertebrates (Horvath 2013), and there is also some evidence among invertebrates (Lian et al. 2015). Meanwhile, several studies have noted considerable variance in methylation patterns that is incompletely explained by genetics and known environmental conditions (Dimond et al. 2017; Durante et al. 2019; Rondon et al. 2017). However, the significant convergence of colony methylation profiles after a year in the common garden together suggest that methylation was in fact responding to, and reflective of, the common garden environment.

Although epigenetic variation among colonies was considerably smaller than genetic variation, there was a weak but significant positive correlation between these two variables, suggesting that methylation patterns are at least partially heritable. A similar relationship was observed among branching *Porites* spp. by Dimond et al. (2017), and additional evidence for heritability of methylation in corals has been reported by several other recent studies (Dixon et al. 2018; Durante et al. 2018; Liew et al. 2018a). While a full appraisal of the potential for methylation to be involved in transgenerational acclimative responses in corals awaits further study, the reports to date are promising.

Lack of an annotated *P. astreoides* genome limited the scope of our analysis and the inferences we were able to draw regarding potential functional implications of changes in methylation, however, a handful of loci exhibiting consistent differences in methylation provide some clues. Most of these loci were apparently associated with coding sequences, which is consistent with gene body methylation as the primary form of methylation among invertebrates (Sarda et al. 2012). One locus was associated with a sequence coding for a calcium-independent protein kinase C-like. Calcium-independent protein kinase C is involved in intracellular signaling, a biological process that is typically associated with the hypomethylated fraction of the genome (Dimond and Roberts 2016). By contrast with hypermethylated housekeeping genes that tend to exhibit consistent expression across conditions and tissues, hypomethylated genes such as those involved in cell-cell signaling are characterized by their inducibility in response to environmental change (Dimond and Roberts 2016).

A gene encoding a putative tax1-binding protein homolog was another differentially methylated locus. These proteins are involved in negative regulation of apoptotic processes via negative regulation of NF-κB transcription factor activity. Interestingly, deregulation of host NF-κB is associated with dinoflagellate symbiosis in cnidarians (Mansfield et al. 2017); loss of symbionts (bleaching) is associated with elevated levels of NF-κB. Perhaps differential methylation of a tax1-binding protein gene was associated with symbiotic homeostatic maintenance in the new environment.

Another locus resembled a gene encoding a U5 small nuclear ribonucleoprotein 200 kDa helicase, which is involved in mRNA splicing via its role in the spliceosome. Epigenetic factors are widely implicated in alternative mRNA splicing, which is a major source of protein diversity, and hence, phenotypic variation (Luco et al. 2011). DNA methylation itself has been identified as a modulator of alternative splicing (Lev Maor et al. 2015).

The last mRNA-associated locus coded for a putative FAM98A protein. These proteins have numerous associated biological processes, including positive regulation of cell proliferation and gene expression. However, the most intriguing function involves protein methylation. A study of human colorectal cancer found that FAM98A was required for expression of an arginine methyltransferase (Akter et al. 2017). Protein methylation has been widely studied in histones, and there is ample evidence for epigenetic crosstalk between methylated DNA and methylated histones (Du et al. 2015).

Further evidence of epigenetic crosstalk in the loci responsive to transplantation is suggested by two loci associated with putative non-coding RNAs (ncRNAs). Along with DNA methylation and histone modifications, ncRNAs are considered part of the epigenetic machinery. ncRNAs are considerably more abundant than mRNAs and play numerous roles, particularly in processes such as post-transcriptional mRNA silencing (Kornienko et al. 2013). In some cases, they are involved in directing DNA and histone methylation patterns (Kornienko et al. 2013; Miska and Ferguson-Smith 2016).

### Conclusion

This work shows that DNA methylation is an environmentally responsive epigenetic process that is reflective of the environment, and is consistent with its putative role in acclimatization. We were able to detect subtle changes in *P. astreoides* methylation associated with experimental transplantation, as well as evidence for heritability of methylation patterns. Loci responding to transplantation were associated with signaling, apoptosis, gene regulation and epigenetic crosstalk, yet much remains to be learned about the function of methylation changes in these differentially methylated genes. This study helps set the stage for further work on both the functional genomics and molecular ecology of acclimatization processes in reef corals.

